# Functional annotation of rare structural variation in the human brain

**DOI:** 10.1101/711754

**Authors:** Lide Han, Xuefang Zhao, Mary Lauren Benton, Thaneer Perumal, Ryan L. Collins, Gabriel E. Hoffman, Jessica S. Johnson, Laura Sloofman, Harold Z. Wang, CommonMind Consortium, Kristen J. Brennand, Harrison Brand, Solveig K. Sieberts, Stefano Marenco, Mette A. Peters, Barbara K. Lipska, Panos Roussos, John A. Capra, Michael Talkowski, Douglas M. Ruderfer

## Abstract

Structural variants (SVs) contribute substantially to risk of many brain related disorders including autism and schizophrenia. However, annotating the potential contribution of SVs to disease remains a major challenge. Here, we integrated high resolution SV calling from genome-sequencing in 755 human post-mortem brains with dorsal lateral prefrontal cortex RNA-sequencing from a subset of 629 samples to quantify the dosage and regulatory effects of SVs. We show that genic (p = 5.44×10^−9^) and regulatory SVs (enhancer p = 3.22×10^−23^, CTCF p = 3.86×10^−18^) are present at significantly lower frequencies than intergenic SVs after correcting for SV length. Copy number variants (CNVs)—deletions and duplications—exhibit a significant quantitative and directional relationship between the proportion of genic and regulatory content altered and gene expression, and the size of the effect is inversely correlated with the loss-of-function intolerance of the gene. We trained a joint linear model that leverages genic and regulatory annotations to predict expression effects of rare CNVs in independent samples (R^2^ = 0.21-0.41). We further developed a regulatory disruption score for each CNV that aggregates the predicted expression across all affected genes weighted by the genes’ intolerance score and applied it to an independent set of SVs from 14,891 genome-sequenced individuals. Pathogenic deletions implicated in neurodevelopmental disorders by ClinGen had significantly more extreme regulatory disruption scores than the rest of the SVs. Rank ordering based on the most extreme regulatory disruption scores prioritized pathogenic deletions that would not have been prioritized by frequency or length alone. This work points to the deleteriousness of regulatory SVs, particularly those altering CTCF sites. We further provide a simple approach for functionally annotating the regulatory effects of SVs in the human brain that has potential to be useful in larger SV studies and should improve as more regulatory annotation data is generated.

## Introduction

Structural variants (SVs) are a common and complex form of genetic variation that contribute substantially to phenotypic diversity and disease^1–3^. This contribution is particularly notable in brain related disorders and traits such as schizophrenia, autism spectrum disorder (ASD) and cognition^4–8^. The advent of short-read genome sequencing has facilitated SV detection at nucleotide resolution and enabled the generation of large-scale reference studies^2,9,10^. Despite this progress, we still have a limited understanding of the functional impact of these variants, particularly for those seen infrequently in populations. Developing approaches to infer the functional consequences of SVs in the brain could have a profound impact on interpretation of genetic risk of complex brain disorders.

RNA-sequencing enables accurate measurement of transcription genome-wide^11^ and can thus facilitate a direct assessment of functional changes driven by genetic variants including single nucleotide variants (SNVs) and SVs^12–15^. Previous work using earlier technologies has demonstrated that SVs have profound effects on expression with estimates of large copy number variants (CNVs) alone explaining 18% of variation in gene expression in cell lines^19^. Most work in this area has focused on common variants for which there is statistical power to identify direct association of a variant with expression of a gene^16^. However, recent analyses have suggested a substantial regulatory role for rare variants by identifying an enrichment of expression outliers in individuals harboring such variation^17,18^. In a small number of examples, rare SVs have shown the potential to alter the expression of genes both within and outside the SV locus with disease relevant phenotypic consequences. Such SVs often alter the regulatory landscape directly or through positional effects that change the threedimensional structure of the genome^19–23^. For example, the expression of *PLP1* is regulated by a downstream duplication and is associated with spastic paraplegia type 2 with axonal neuropathy^22^. To date, few samples have thus far been able to leverage both comprehensive SV detection from genome-sequencing and RNA-sequencing to explore the effects of rare SVs on expression genome-wide. Despite the importance of rare SVs in brain-related disorders and the tissue specific nature of transcriptional regulation, efforts to understand the functional consequences of rare SVs in the brain have been impeded by the challenge in acquiring enough post-mortem brain samples to be well-powered to quantify the dosage and regulatory effects of SVs

The CommonMind Consortium (CMC; www.synpase.org/CMC) is a large collection of collaborating brain banks that includes over 1,000 samples. Here, we leveraged newly generated genome-sequencing data integrated with RNA-sequencing data in 629 samples that enabled us to directly study the effects of rare SVs on expression in the brain. We show that SVs affecting regulatory elements are at significantly lower variant frequencies than expected, suggesting their potential to be deleterious. We also provide a quantitative characterization of the effects of SVs altering different regulatory elements have on expression. These results show that most complete gene deletions and duplications do not result in expression outliers and that genic intolerance to variation informs their functional impact. Finally, we built a model to infer the expression effects of SVs and used it to calculate a cumulative measure of regulatory disruption of an SV across all genes. When applied to a large independent SV reference data set^9,24^, the regulatory disruption score improves prioritization of pathogenic deletions beyond the common practice of considering frequency and SV length. Altogether, this work advances our understanding of the transcriptional consequences of SVs in the human brain and provides a framework for functionally annotating these variants to aid in disease studies.

## Methods

### Cohorts

Samples were included from two different cohorts. The CMC study is a combined collection of brain tissues from the Mount Sinai NIH Brain Bank and Tissue Repository (n=127), The University of Pennsylvania Brain Bank of Psychiatric Illnesses and Alzheimer’s Disease Core Center (n=62) and The University of Pittsburgh NIH NeuroBioBank Brain and Tissue Repository (n=139). The CMC_HBCC study includes brain samples from the NIMH Human Brain Collection Core (n=445).

### DNA Isolation

DNA for all 773 samples was isolated from approximately 50 mg dry homogenized tissue from the dorsal lateral prefrontal cortex (DLPFC). All tissue samples had corresponding tissue samples that were isolated for RNAseq. All DNA isolation was done using the Qiagen DNeasy Blood and Tissue Kit (Cat#69506) according to manufacturer’s protocol. DNA yield and genomic quality number (GQN) was quantified using Thermo Scientific’s NanoDrop and the Fragment Analyzer Automated CE System (Advanced Analytical). 96 samples had a GQN <4, but were not excluded from genome-sequencing. The mean yield was 9.9 ug (SD = 10.4) and the mean GQN was 5.6 (SD = 1.47).

### DNA Library Preparation

All samples were submitted to the New York Genome Center for genome-sequencing, where they were prepared for sequencing in randomized batches of 95. The sequencing libraries were prepared using the Illumina PCR-free DNA sample preparation Kit. The insert size and DNA concentration of the sequencing library was determined on Fragment Analyzer Automated CE System (Advanced Analytical) and Quant-iT PicoGreen (ThermoFisher) respectively. A quantitative PCR assay (KAPA), with primers specific to the adapter sequence, was used to determine the yield and efficiency of the adaptor ligation process.

### Genome-sequencing library preparation and sequencing

Libraries for genome sequencing were generated from 100 ng of genomic DNA using the Illumina TruSeq Nano DNA HT sample preparation kit. Genomic DNA were sheared using the Covaris sonicator (adaptive focused acoustics), followed by end-repair, bead-based size selection, A-tailing, barcoded-adaptor ligation followed by PCR amplification. Final libraries were evaluated using qPCR, picogreen and Fragment analyzer. Libraries were sequenced on a 2×150 bp run of a HiSeq X instrument.

### Genome-sequencing pipeline

Paired-end 150 bp reads were aligned to the GRCh37 human reference using the Burrows-Wheeler Aligner (BWA-MEM v0.78) and processed using the best-practices pipeline that includes marking of duplicate reads by the use of Picard tools (v1.83, http://picard.sourceforge.net), realignment around indels, and base recalibration via Genome Analysis Toolkit (GATK v3.2.2). Variants were called using GATK HaplotypeCaller, which generates a single-sample GVCF. To improve variant call accuracy, multiple single-sample GVCF files were jointly genotyped using GATK GenotypeGVCFs, which generated a multi-sample VCF. Variant Quality Score Recalibration (VQSR) was performed on the multi-sample VCF, which added quality metrics to each variant that can be used in downstream variant filtering.

### Structural Variant Discovery

SVs were detected using a discovery pipeline previously described^25^ that relies upon an ensemble of SV detection algorithms to maximize sensitivity, followed by a series of filtering modules to control the overall false discovery rate and refine variant predictions. In brief:

1. **Raw SV calls collection:** Five algorithms that used discordant pair-end reads (PE) and split reads (SR) to predict SVs, i.e. Delly^26^ (v0.7.5), Lumpy^27^ (v0.2.13), Manta^28^ (v1.01), Wham^29^ (v1.7.0) and MELT^30^ (v2.1.4), were executed in per-sample mode with their default parameter. A series of read depth-based (RD) algorithms were also applied for copy number variant (CNV) detection, including CNVnator^31^ (v0.3.2), GenomeSTRiP^32^ (v2), and a custom version of cn.MOPS^25^. These algorithms were applied to male and female samples separately, each in ~100-sample batches. For each batch, we composed a coverage matrix across all samples at 300 bp and 1 kb bin sizes across each chromosome with N-masked bases excluded, then applied cn.MOPS, split raw calls per sample, segregated calls into deletions (copy number < 2) and duplications (copy number > 2) and merged the 300 bp and 1 kb resolution variant predictions per sample per CNV class using *BEDTools merge*.
2. **Aberrant alignment signature collection:** We collected discordant PE and SR evidence through *svtk collectpesr*, RD evidence through *svtk bincov* with N-masked regions excluded (https://github.com/talkowski-lab/CommonMind-SV/blob/master/files/h37.Nmask.bed), and B allele frequency (BAF) evidence from *GATK HaplotypeCaller-generated* VCFs using a custom script (*vcf2baf*, https://github.com/talkowski-lab/CommonMind-SV/blob/master/scripts/vcf2baf.sh). Following evidence collection per sample, we constructed PE, SR, RD, and BAF matrices merged across each phase of sequencing that included 327 (SKL_10073), 326 (SKL_11154) and 119 (SKL_11694) samples respectively through customized scripts (https://github.com/talkowski-lab/CommonMind-SV/tree/master/Step1b_EvidenceCollection).
3. **SV integration and refinement:** SV calls detected by each algorithm described above were integrated and calibrated through a series of filtering modules to control the overall false discovery rate (FDR) and refine variant predictions. Raw outputs from each algorithm were clustered across all samples for each of three sequencing phases (327 samples in SKL_10073; 326 samples in SKL_11154; 119 samples in SKL_11694). Once clustered across samples, the integrated call set was filtered through a random-forest module that tests for statistically significant differences between samples with and without each SV based on four semi-orthogonal signatures: PE, SR, RD, and BAF. Finally, filtered, high-quality SV calls were integrated across all three sequencing phases, whereupon all breakpoints were genotyped, followed by alternate allele structure resolution, complex SV classification, other variant refinements, and gene annotation. The filtering module is adaptable to multiple input algorithms, and this same pipeline has been applied to WGS data in ASD families^25^ and population variation datasets^9^.

### SV dataset description

We successfully applied all SV discovery algorithms on 772 / 773 (99.9%) of the CMC samples, with one failed sample (MSSM-DNA-PFC-375). All 772 samples were included in the SV integration pipeline with SVs assigned in the final call set. The final analyses yielded 125,260 SVs, including 62,948 deletions, 30,547 duplications, 31,155 insertions, 268 simple inversions, 341 complex SVs and 1 reciprocal translocation. On average, 6,220 SVs were identified per sample, consisting of 3,579 deletions, 755 duplications, 1,839 insertions, 15 inversions and 14 complex SVs. For the insertions, 1,146 were further classified as mobile elements insertions (MEI), including 1,005 Alu, 92 LINE1 and 49 SVA variants. The number of SVs distributed proportionately by read depth among the phases and matched expected demographic history (e.g. 1,421 more SVs were detected on average for African-American individuals than all other populations).

### Identification and removal of SV outliers

We carefully examined the set of 772 individuals with SV calls for technical outliers (e.g. related to genome-sequencing generation or biological processes associated with DNA extraction of the post-mortem samples). We removed one individual for presence of an abnormal sex chromosome (XXY) which had been previously noted^33^. We further identified 16 samples that represented CNV outliers resulting from anomalous read dosage as calculated by our dosage scoring metric^9^. Individuals were removed if they carried too many or too few CNVs, defined by 3*IQR (interquartile range) of CNVs per individual or genomic content that they alter. After outlier exclusion, we retained 755 (97.8%) of samples with SV data.

### RNA-sequencing pipeline

The processing of the RNA-sequencing for these samples has been previously described ^34^, however we reiterate this process below for reading convenience. Samples were processed separately by cohort: CMC and CMC_HBCC.

### RNA-sequencing Re-alignment

RNA-sequencing reads were aligned to GRCh37 with STAR v2.4.0g1^51^ from the original FASTQ files. Uniquely mapping reads overlapping genes were counted with featureCounts v1.5.2^52^ using annotations from ENSEMBL v75.

### RNA-sequencing Normalization

To account for differences between samples, studies, experimental batch effects and unwanted RNA-sequencing specific technical variations, we performed library normalization and covariate adjustments using fixed/mixed effects modeling. The workflow consisted of following steps:

1. ***Gene filtering:*** Out of ~56K aligned and quantified genes only genes showing at least modest expression were used in this analysis. Genes that were expressed more than 1 CPM (read Counts Per Million total reads) in at least 50% of samples in each study were retained for analysis. Additionally, genes with available gene length and percentage GC content from BioMart December 2016 archive were subselected from the above list. This resulted in approximately 14K to 16K genes in each batch.
2. ***Calculation of normalized expression values:*** Sequencing reads were then normalized in two steps. First, conditional quantile normalization (CQN)^53^ was applied to account for variations in gene length and GC content. In the second step, the confidence of sampling abundance was estimated using a weighted linear model using voom-limma package in bioconductor^54,55^. The normalized observed read counts, along with the corresponding weights, were used in the following steps.
3. ***Outlier detection:*** Based on normalized log2(CPM) of expression values, outlier samples were detected using principal component analysis (PCA)^56,57^ and hierarchical clustering. Samples identified as outliers using both the above methods were removed from further analysis.
4. ***Covariate identification:*** Normalized log2(CPM) counts were then explored to determine which known covariates (both biological and technical) should be adjusted. For the CMC study, we used a stepwise (weighted) fixed/mixed effect regression modeling approach to select the relevant covariates having a significant association with gene expression. Here, covariates were sequentially added to the model if they were significantly associated with any of the top principal components, explaining more than 1% of variance of expression residuals. For CMC_HBCC, we used a model selection based on Bayesian information criteria (BIC) to identify the covariates that improve the model in a greater number of genes than making it worse.
5. ***SVA adjustments:*** After identifying the relevant known confounders, hidden-confounders were identified using the Surrogate Variable Analysis (SVA)^58^. We used a similar approach as previously defined^49^ to find the number of surrogate variables, which is more conservative than the default method provided by the SVA package in R^59^. The basic idea of this approach is that for an eigenvector decomposition of permuted residuals each eigenvalue should explain an equal amount of the variation. By the nature of eigenvalues, however, there will always be at least one that exceeds the expected value. Thus, from a series of 100 permutations of residuals (white noise) we identified the number of covariates to include. We applied the “irw” (iterative re-weighting) version of SVA to the normalized gene expression matrix, along with the covariate model described above to obtain residual gene expression.
6. ***Covariate adjustments:*** We performed a variant of fixed/mixed effect linear regression, choosing mixed-effect models when multiple samples, were available per individual, as shown here: gene expression ~ Diagnosis + Sex + covariates + (1|Donor), where each gene in linearly regressed independently on Diagnosis, identified covariates and donor (individual) information as random effect. Observation weights (if any) were calculated using the voom-limma^54,55^ pipeline, which has a net effect of up-weighting observations with inferred higher precision in the linear model fitting process to adjust for the meanvariance relationship in RNA-sequencing data. The Diagnosis component was then added back to the residuals to generate covariate-adjusted expression.

All these workflows were applied separately for each cohort. For CMC_HBCC, samples with age < 18 were excluded prior to analysis.

### Ensuring sample consistency between genome-sequencing and RNA-sequencing

To infer effects on expression from SVs, we had to ensure the genome-sequencing and RNA-sequencing data were from the same individual. To do so we used variant calling data from both platforms. We removed SNVs with missing rate ≥ 0.05 and restricted only to biallelic variants. Upon merging the genotypes from genome-sequencing and RNA-sequencing we calculated genome-wide relatedness from estimates of identity-by-descent using Plink^60^ across all cross-platform pairs of samples. For each genome-sequencing sample we identified the appropriate matching RNA-sequencing sample requiring both near complete relatedness (Pihat > 0.8) and no other sample with high relatedness. Across both cohorts 622/632 (98%) samples matched the expected pair and 10 samples had to be corrected.

### Genomic annotation sources

All data were downloaded in the GRCh37/hg19 build of the human genome. We used TSS definitions from Ensembl v75. To map regions of open chromatin, we used a set of DNase hypersensitive sites (DHSs) downloaded from Roadmap Epigenomics^61^. We mapped the three-dimensional chromatin architecture using TAD domains identified by PsychENCODE from Hi-C contact matrices with 40 kb resolution in the prefrontal cortex (PFC, n=2,735)^62^. As a proxy for TAD boundaries or other insulated regions, we used a set of CTCF binding sites from ChIP-seq data downloaded from ENCODE in brain-relevant cell types^63^. We merged overlapping CTCF peaks from each tissue into a single consensus region (n=100,894).

### Cis-regulatory element annotations

We downloaded PFC enhancer annotations (n=79,056) from the PsychENCODE project^62^. These were generated by overlapping cross-tissue DNase-seq and ATAC-seq assay information with H3K27ac ChIP-seq peaks. Regions overlapping H3K4me3 peaks and within 2 kb of a TSS were excluded from the set of putative enhancers. All ChIP-seq, ATAC-seq, and DNase-seq data were filtered to include only high-signal peaks with a z-score greater than 1.64. We also downloaded the high confidence set of enhancer annotations (n=18,212) which, in addition to the criteria above, require high PFC H3K27ac ChIP-seq signal (z-score > 1.64) in both the PsychENCODE and Roadmap Epigenomics experiments. We generated a set of promoter annotations by using 2 kb windows upstream from each TSS (n=57,773). We intersected these 2 kb windows with PFC H3K27ac from PsychENCODE and PFC H3K4me3 from Roadmap Epigenomics to create a set of high confidence promoters (n=5,73 6)^61,62^. As in the enhancer definition, the H3K27ac and H3K4me3 ChIP-seq data included only high signal peaks with a z-score > 1.64.

## Results

### Evidence for selection against SVs affecting transcriptional regulation

The SV detection pipeline identified 116,471 high-quality variants across 755 individuals. The final set of SVs predominantly consisted of CNVs (73%) and mobile element insertions (18%). The vast majority of SVs were small and rare (Figure 1, Supplementary Table 1). The average length of SVs in this dataset was 7,053 bp (median = 280 bp), with 83% of variants less than 1 kb. We next identified a subset of large (> 1 kb) and rare SVs (observed in < 1% of individuals), representing 20,001 variants, 91% of which were CNVs given that most mobile elements are small. On average, individuals carried 75.8 rare, large SVs including 43.6 deletions and 25.7 duplications. These numbers differed by ancestry; individuals with African ancestry (mean = 121.9 SVs) carried substantially more rare SVs than individuals with European (mean = 51 SVs) or other ancestries, as expected.

**Figure 1.**
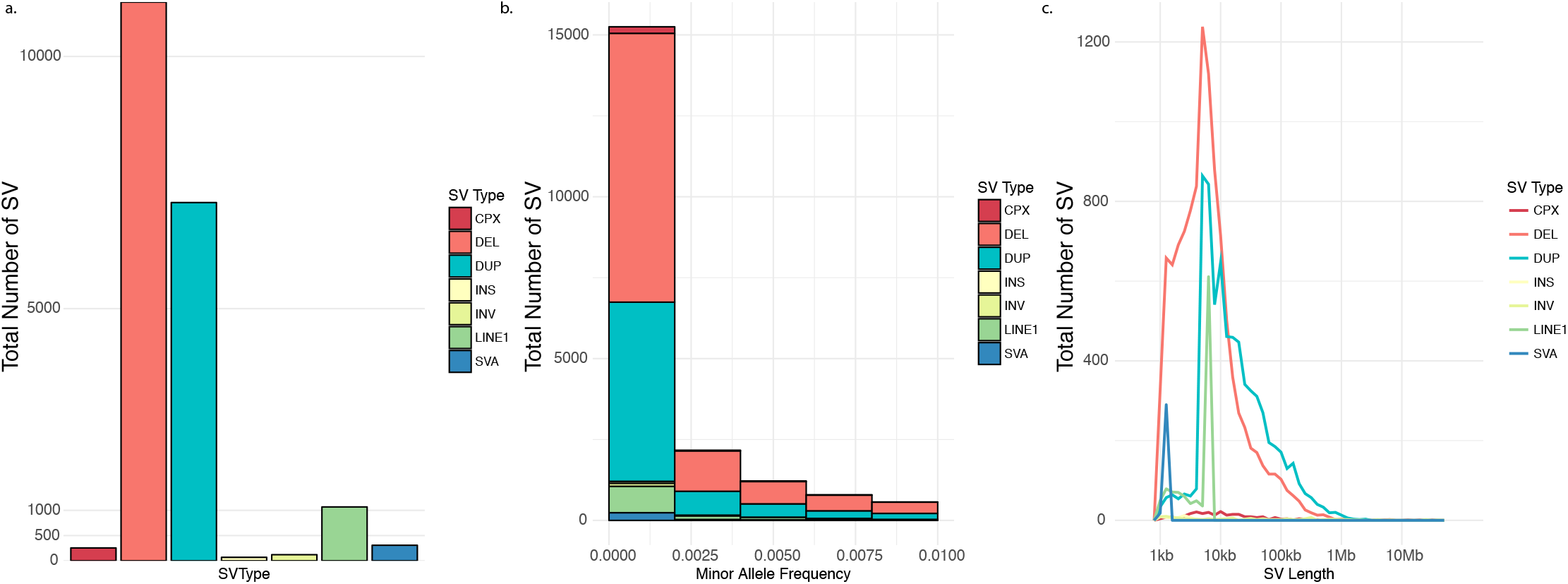
Characterization of high confidence (>1 kb) and rare (<1%) SV dataset stratified by a) type of SV, b) minor allele frequency and c) length (log10-scaled) colored by type of SV. SV Types include complex (CPX), deletion (DEL), duplication (DUP), insertion (INS), inversion (INV), long interspersed nuclear element-1 (LINE1), combined set of mobile elements (SVA) including short interspersed nuclear elements, variable number tandem repeat and Alu.

We next sought to characterize how frequently SVs putatively alter gene dosage based on overlap with genes or regulatory elements. We defined a set of regulatory elements that included CTCF sites (n=100,894), enhancers (n=79,056) derived exclusively from brain tissue (see Methods) and promoters (2 kb upstream of the transcription start site [TSS]). Genes were defined as those in Ensembl v75 (n=57,773) and where noted we split protein-coding genes from others which we label broadly as “other transcribed products.” For comparison, we defined two “non-functional” categories of SVs: intronic and intergenic SVs lacking overlap with any annotation. Despite this annotation, these non-functional SV categories will include some proportion of SVs altering functional elements that were either not included, or not yet identified, which should make our comparisons conservative. The allele frequency (AF) of SVs affecting protein-coding genes (AF = 0.00167, p = 5.44×10^−9^), enhancers (AF = 0.0012, p = 3.22×10^−23^) and CTCF sites (AF = 0.00156, p = 3.86×10^−18^) were significantly lower and singleton proportions were significantly higher than intergenic SVs (mean AF = 0.00193) after matching for SV length (Table 1a, Wilcoxon test).

**Table 1.** Frequency measures (allele frequency [AF] and singleton proportion) of all SVs a) overlapping a particular annotation and b) uniquely overlapping a single annotation. Both are then matched 1 to 1 to intergenic SVs based on length within 500 bp and tested for frequency differences using Wilcoxon Rank Sum test. Table 1c presents AF and singleton proportion for all SVs overlapping a particular annotation in gnomAD. Red text indicates p-value < 0.05.

| Matched to intergenic SV by length |  |  |  |  |  |  |  |  |  |  |  |  |
| --- | --- | --- | --- | --- | --- | --- | --- | --- | --- | --- | --- | --- |
| a. | Annot | N | Length | AF | Singleton Proportion | N | Length | Length (intergenic) | p-value (length) | AF | AF (intergenic) | p-value |
|  | coding | 4145 | 15683 | 0.00163 | 0.644 | 2864 | 15401.6 | 15350.9 | 0.99 | 0.00167 | 0.00192 | **5.4E-09** |
|  | intronic | 5081 | 5000 | 0.00195 | 0.544 | 5048 | 7185.4 | 7240.0 | 0.34 | 0.00196 | 0.00204 | **3.4E-02** |
|  | enhancer | 1366 | 16816.5 | 0.00118 | 0.774 | 1012 | 20092.2 | 20063.7 | 0.98 | 0.00120 | 0.00188 | **3.2E-23** |
|  | promoter | 1471 | 37221 | 0.00178 | 0.580 | 946 | 25721.4 | 25704.2 | 0.99 | 0.00186 | 0.00189 | 8.8E-01 |
|  | ctcf | 4481 | 21641 | 0.00146 | 0.693 | 2706 | 15815.2 | 15748.0 | 0.84 | 0.00156 | 0.00191 | **3.9E-18** |
|  | intergenic | 7220 | 5450 | 0.00202 | 0.540 |  |  |  |  |  |  |  |
| Matched to intergenic SV by length |  |  |  |  |  |  |  |  |  |  |  |  |
| b. | Annot | N | Length | AF | Singleton Proportion | N | Length | Length (intergenic) | p-value (length) | AF | AF (intergenic) | p-value |
|  | coding | 1223 | 7000 | 0.00187 | 0.597 | 1215 | 10835.7 | 10813.5 | 0.93 | 0.0019 | 0.0020 | **2.0E-02** |
|  | intronic | 3723 | 3988 | 0.00200 | 0.528 | 3719 | 5476.8 | 5543.5 | 0.26 | 0.0020 | 0.0021 | 7.2E-01 |
|  | enhancer | 213 | 5500 | 0.00182 | 0.610 | 214 | 7960.2 | 7982.8 | 0.97 | 0.0018 | 0.0020 | 3.5E-01 |
|  | promoter | 91 | 3019 | 0.00231 | 0.462 | 92 | 3979.0 | 3950.4 | 0.85 | 0.0023 | 0.0021 | 3.5E-01 |
|  | ctcf | 1332 | 9000 | 0.00165 | 0.630 | 1267 | 16007.5 | 15967.1 | 0.93 | 0.0017 | 0.0020 | **1.0E-04** |
|  | intergenic | 7220 | 5450 | 0.00202 | 0.540 |  |  |  |  |  |  |  |
| c. | Annot | N | Length | AF | Singleton Proportion |  |  |  |  |  |  |  |
|  | coding | 19577 | 14881 | 0.00045 | 0.574 |  |  |  |  |  |  |  |
|  | enhancer | 7653 | 24000 | 0.00029 | 0.631 |  |  |  |  |  |  |  |
|  | promoter | 22378 | 21000 | 0.00053 | 0.524 |  |  |  |  |  |  |  |
|  | ctcf | 25364 | 21500 | 0.00035 | 0.584 |  |  |  |  |  |  |  |
|  | other | 53046 | 4229.5 | 0.00046 | 0.523 |  |  |  |  |  |  |  |

To explore the contributions of different functional elements to this result, we stratified SVs based on the specific annotations (*e.g.*, coding and enhancer, Supplementary Figure 1) to isolate those that alter combinations and those that uniquely alter a single annotation (Table 1b). We identified a significant negative correlation between the total number of annotations and minor allele frequency indicating that SVs with more potential to alter dosage are less likely to be tolerated (Supplementary Figure 2). Further, we show that SVs exclusively affecting CTCF sites (AF = 0.0017, p = 0.0001) had the lowest frequency and the most significant result when compared to intergenic variation at a level comparable to SVs that only affected protein-coding genes (AF = 0.0019, p = 0.02). These results are consistent across SV type and this difference in allele frequency is seen when performing the same annotation of the gnomAD SV dataset of ~15k samples called from genome-sequencing using the same pipeline^9^ (Table 1c). These results suggest a similarly strong selection against SVs that alter CTCF sites and protein-coding genes, as presented previously^64^. Since not all genes are equally sensitive to dosage changes, we split protein-coding genes into two sets based on whether they were more constrained on average as defined from gnomAD loss-of-function observed/expected score < 0.49 and recalculated AF and singleton proportion. SVs affecting the genes in the more intolerant half had lower frequencies (AF = 0.00138) than those affecting CTCF sites (AF = 0.00146).

### Transcriptional consequences of regulatory SVs

Among the samples with genome-sequencing, 629 individuals had RNA-sequencing data from the dorsal lateral pre-frontal cortex (DLPFC). RNA-sequencing was done across two cohorts (CMC and CMC_HBCC), results were consistent across cohorts as shown in many instances. To quantify the transcriptional consequence of an SV, we defined expression in two ways. First, we calculated relative expression as the average expression of carriers divided by non-carriers. Second, we calculated z-scores using only non-carriers for calculating the mean and standard deviation. We use both measures throughout, relying on relative expression in certain cases for interpretation but preferring z-scores for their statistical properties.

We expect complete deletion or duplication of a gene to result in a 50% decrease or increase in expression, respectively. Relative expression calculated using read counts per million total reads (CPM) demonstrated the expected 50% decrease or increase from full gene deletions or duplications, on average (Supplementary Figure 3). Deletions fit this expectation better than duplications, suggesting more variability among duplication calls and/or their functional effects. Normalization and linear covariate adjustment, which is necessary to account for confounders and batch effects (see Methods) alters the relative difference in expression among carriers to be closer to 25% while also reducing the variance, enabling clear demonstration of expression differences among individuals carrying full gene deletions or duplications (Supplementary Figure 3). In general, expression was substantially lower across the full set of deletions and higher across the full set of duplications affecting genes (Figure 2), and these results were consistent across RNA-sequencing cohorts (Supplementary Figure 4). Not surprisingly, other classes of SVs did not show a directional expression difference.

**Figure 2.**
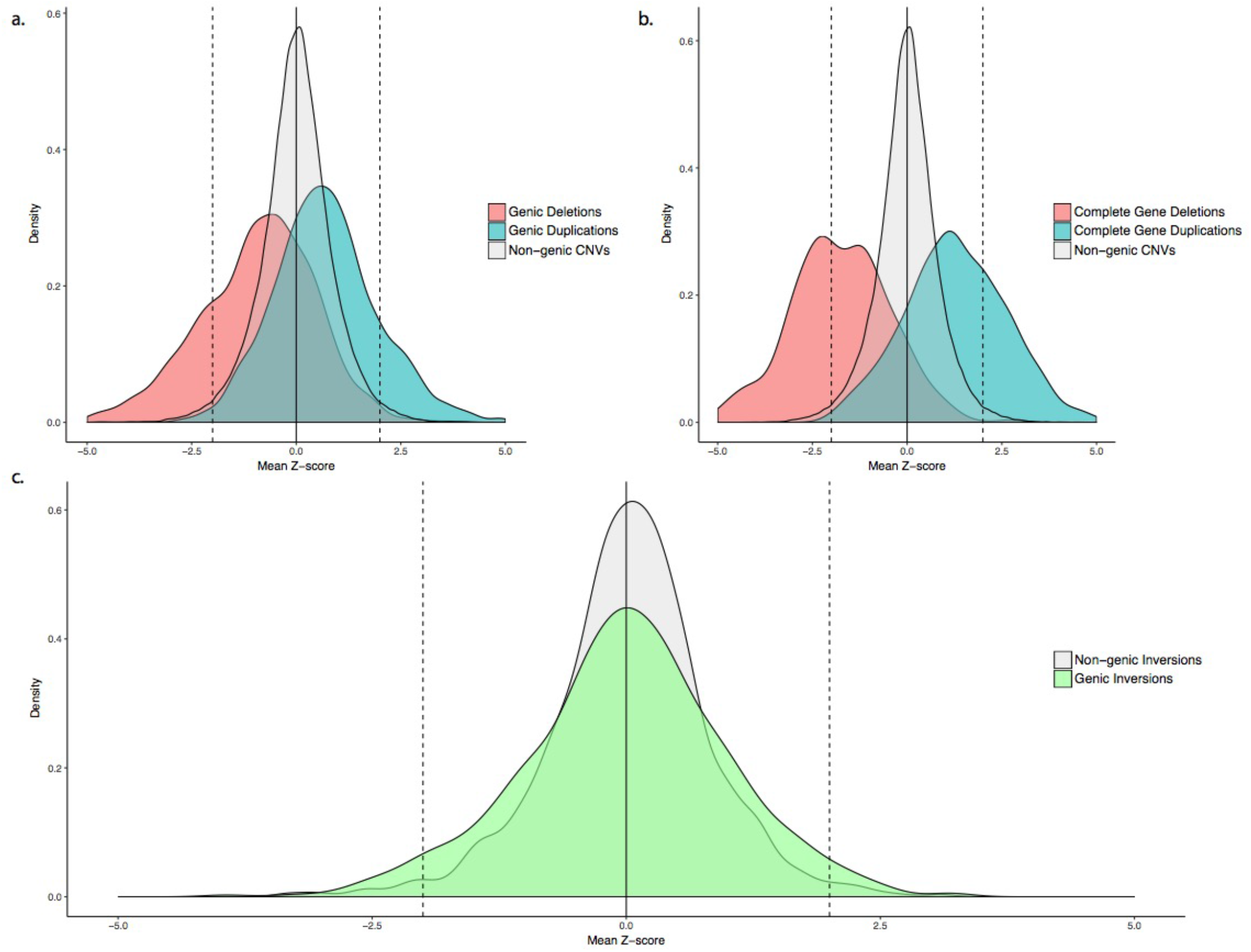
Expression presented as a z-score for a) all CNV that overlap any proportion of the exonic sequence of a gene, b) CNV that delete or duplicate 100% of the exonic sequence of a gene and c) all inversions with any gene overlap (green) compared to all other SV (grey). Deletions are red, duplications are blue. The dashed lines are located at z-score of 2 and −2.

While there is an expectation for the effect of full gene deletions and duplications, the effects that other SVs may have on expression are not obvious. We identified a relationship between proportion of exonic sequence deleted/duplicated and expression (Supplementary Figure 5) where the more exonic sequence deleted or duplicated the more extreme the expression difference. We defined expression outliers as those with z-scores greater than 2 or less than −2, and considered all pairs of SV and genes within 1 Mb of the SV. After Bonferroni correcting for 36 tests (p < 0.0014), we identified significant excess of positive expression outliers for genic duplications (14.6%, p = 8.4x-10^−158^, Fisher’s exact test) and significant excess of negative expression outliers for genic deletions (22.8%, p = 8.4×10^−163^) when compared within the same CNV type but not affecting proteincoding genes (Table 2). These results remained consistent whether we tested protein-coding genes or other transcribed gene products (Table 2). Among the other SV classes, only inversions provided a large enough set to test, and we identified a significant excess of expression outliers among genic inversions when considering outliers in both directions (6.8%, p = 1.2×10^−4^, Table 2), with a slightly larger contribution from positive expression outliers (4.1%, p = 1.1×10^−3^) than negative expression outliers (2.7%, p = 3.2×10^−2^)

**Table 2.**
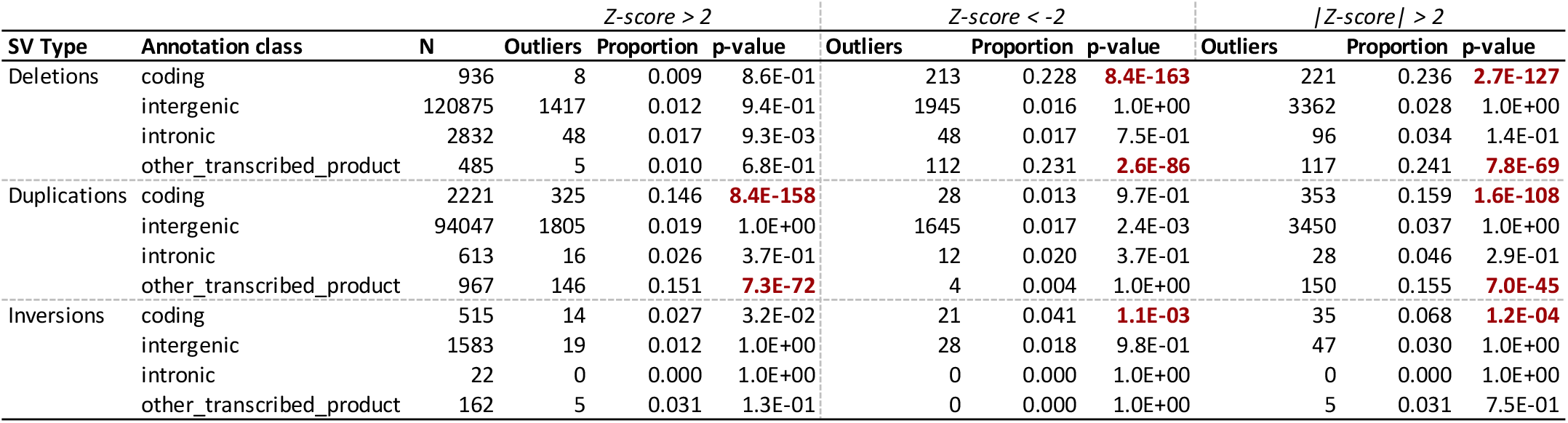
Number and proportion of expression outliers by SV type and annotation. P-values are from Fisher’s Exact test comparing SVs in annotation class to others within SV type. Red text indicates significance after Bonferroni multiple test correction for 36 tests (p < 0.0014).

Our data show an enrichment of expression outliers among genic SVs. However, we emphasize that the impact of structural rearrangement on expression is non-uniform and more complex than a simple accounting of the presence or absence of an SV. Even in the most extreme case of 100% gene deletion or duplication, the affected gene would only be considered an expression outlier in 47% of deletions and 30% of duplications (Figure 2). Furthermore, across all genic CNVs, only 22.8% of genic deletions and 14.6% of genic duplications result in gene expression outliers. These results demonstrate that the majority of known genic SVs would not be identified if restricted to expression outliers, so in contrast to this thresholding approach, a quantitative approach can more accurately assess the effects that SVs have on gene expression.

Next, we quantified the transcriptional consequences of SVs that affect regulatory elements; this requires determination of a set of genes to analyze for each element. Our definition of promoters was necessarily gene specific; however, for enhancers we explored numerous approaches concluding that genes predicted to be targets of enhancers from Hi-C^65^ was the most interpretable and useful for downstream analyses. Other approaches that we considered including the nearest gene, all genes within a shared topological associating domain (TAD) and all genes within a 1 Mb window showed similar results. We therefore included 90,015 enhancer-gene pairs covering 6,535 genes and 32,803 enhancers predicted from PsychENCODE Hi-C data^65^. To capture the relative contributions of all annotations, we tested the relationship between gene expression z-score and SV annotations with a joint linear model that included exon proportion, promoter proportion, sum proportion of all affected enhancers, whether SV and gene were within the same TAD and SV length. The most significant contributor to expression was the proportion of the exon affected (deletions: beta = −26.925, p = 3.2.×10^−159^; duplications: beta = 21.834, p = 2×10^−105^). Expression was significantly and positively correlated with the proportion of a promoter that was affected by CNVs with deletions leading to lower expression (beta = −3.093, p = 2×10^−3^) and duplications leading to higher expression (beta = 11.324, p = 1×10^−29^). Further, expression was significantly correlated with the cumulative sum of enhancer sequence that was affected by an SV only in duplications, but both deletions and duplications led to decreased expression (deletions: beta = −1.825, p = 0.068; duplications: beta = −5.913, p = 3.4×10^−9^). The presence of the SV and the gene within the same TAD contributed significantly and directionally to expression (deletions: beta = −2.694, p = 7.1×10^−3^; duplications: beta = 4.114, p = 3.9×10^−5^). The effects of these variables on expression were consistent across cohort (Table 3) and while exon proportion provided the strongest contributor, the effects of cis-regulatory elements remained significant in duplications after excluding all genic SVs (Supplementary Table 2).

**Table 3.** Coefficients of linear model to predict expression z-score in a) deletions and b) duplications, across all samples and stratified by cohort. Red cells are those with p-values < 0.05.

| a. | Variable |  |  |  |  | CMC |  |  |  | CMC_HBCC |  |  |  |
| --- | --- | --- | --- | --- | --- | --- | --- | --- | --- | --- | --- | --- | --- |
|  |  | Est. | SE | T | P | Est. | SE | T | P | Est. | SE | T | P |
|  | Exon Proportion | -1.733 | 0.064 | -26.925 | **3.2E-159** | -2.016 | 0.084 | -24.069 | **1.6E-127** | -1.534 | 0.088 | -17.484 | **2.6E-68** |
|  | Enhancer sum | -0.015 | 0.008 | -1.825 | 6.8E-02 | -0.007 | 0.009 | -0.721 | 4.7E-01 | -0.021 | 0.011 | -1.968 | **4.9E-02** |
|  | Promoter proportion | -0.178 | 0.058 | -3.093 | **2.0E-03** | -0.136 | 0.074 | -1.846 | 6.5E-02 | -0.143 | 0.079 | -1.806 | 7.1E-02 |
|  | SV Length | -2.15E-07 | 0.000 | -15.194 | **4.3E-52** | -1.98E-07 | 1.67E-08 | -11.815 | **3.5E-32** | -1.90E-07 | 3.30E-08 | -5.756 | **8.6E-09** |
|  | Within TAD | -0.013 | 0.005 | -2.694 | **7.1E-03** | -0.019 | 0.006 | -2.933 | **3.4E-03** | -0.007 | 0.006 | -1.183 | 2.4E-01 |

| b. | Variable |  |  |  |  | CMC |  |  |  | CMC_HBCC |  |  |  |
| --- | --- | --- | --- | --- | --- | --- | --- | --- | --- | --- | --- | --- | --- |
|  |  | Est. | SE | T | P | Est. | SE | T | P | Est. | SE | T | P |
|  | Exon Proportion | 0.769 | 0.035 | 21.834 | **2.0E-105** | 1.128 | 0.054 | 20.974 | **3.6E-97** | 0.492 | 0.043 | 11.449 | **2.6E-30** |
|  | Enhancer sum | -0.017 | 0.003 | -5.913 | **3.4E-09** | 0.001 | 0.006 | 0.150 | 8.8E-01 | -0.017 | 0.003 | -5.733 | **9.9E-09** |
|  | Promoter proportion | 0.366 | 0.032 | 11.324 | **1.0E-29** | 0.324 | 0.050 | 6.493 | **8.5E-11** | 0.350 | 0.039 | 9.040 | **1.6E-19** |
|  | SV Length | 3.8E-07 | 0.000 | 13.691 | **1.3E-42** | 3.78E-07 | 3.75E-08 | 10.086 | **6.8E-24** | 3.08E-07 | 3.87E-08 | 7.954 | **1.8E-15** |
|  | Within TAD | 0.024 | 0.006 | 4.114 | **3.9E-05** | 0.013 | 0.009 | 1.381 | 1.7E-01 | 0.028 | 0.007 | 3.900 | **9.6E-05** |

### Integrating transcriptional consequences and measures of gene intolerance

To better understand the relationship between our variant annotations in the context of the genes affected, we incorporated two distinct measures of genic intolerance to variation: (1) CNV intolerance defined empirically from exome-sequencing in nearly 60,000 individuals^66^, and (2) a measure of gene intolerance to loss-of-function (LoF) variation generated from a sample of ~141,000 individuals^24^. SVs affecting genes that are more intolerant to variation had smaller expression effects on average (Figure 3). Multiple factors contributed to this pattern. These genes were significantly less likely to have a genic SV (pLoF= 1.01×10^−53^, pCNV = 6.8×10^−39^, Wilcoxon test of gene metric by whether SV is exonic or not) and when they do, they were also significantly likely to alter a smaller proportion of the exonic sequence (pLoF = 1.05×10^−53^, pCNV = 1.28×10^−38^, Spearman correlation test). Notably, these results were driven by CNVs with more significant effects of deletions when using LoF and duplications when using CNV intolerance (Figure 3). Restricting to SVs that only alter regulatory elements and not coding sequence, we identified a significant decrease in the number of enhancers affected by SVs in genes with higher intolerance, although this is only observed for the CNV intolerance metric (pLoF = 0.19, p CNV = 7.7×10^−13^). We did not find any effects from promoter SVs in either metric (pLoF = 0.36, pCNV = 0.09). Combined with the differences seen by CNV type these results may indicate unique properties of these metrics and what they reflect (e.g. haploinsufficiency vs dosage sensitivity). In general, as with single nucleotide variation, genic measures of intolerance should help functionally annotate SVs.

**Figure 3.**
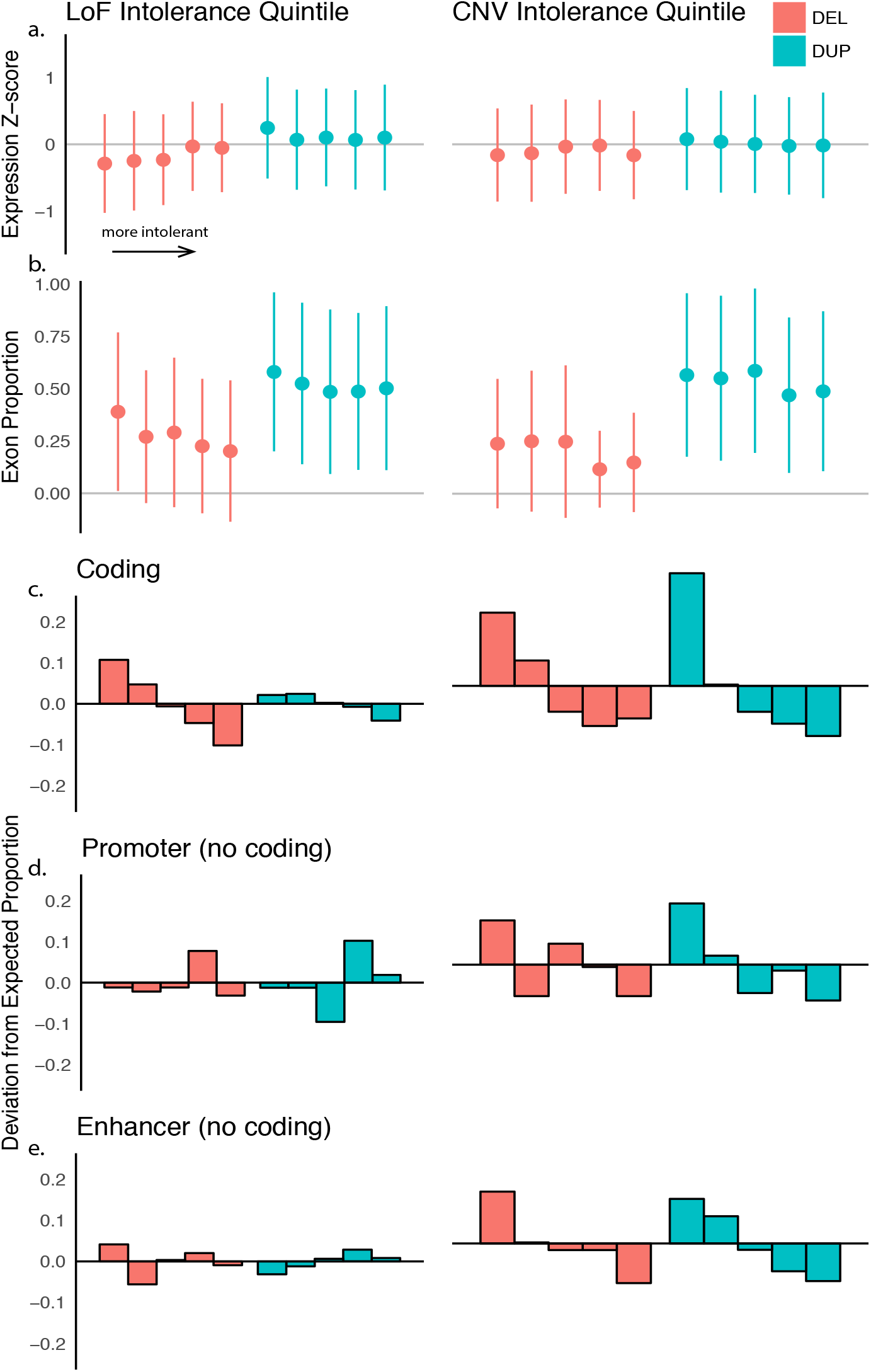
Expression effects of SVs as a product of affected gene intolerance to deleterious variation. Each plot stratifies genes using either the LoF intolerance metric or the CNV intolerance metric into quintiles (20% bins) ordered left to right from least to most intolerant genes and by deletion (red) and duplication (blue). The plots show the effect of this stratification on a) overall expression (z-score) of genic CNV showing mean and standard deviation, b) the proportion of the exonic sequence that is affected showing mean and standard deviation, c) the deviation from the expected 20% of CNV that alter exonic sequence, d) the deviation from expected for non-exonic CNV that alter promoters and e) the deviation from expected for non-exonic CNV that alter enhancers.

### Building a model to annotate SVs from predicted dosage and gene intolerance

Having demonstrated a significant role for SVs in altering expression, we sought to test whether this model could be used to predict expression effects of SVs in independent samples. We split our DLPFC sample by cohort (CMC and CMC_HBCC, see Methods) and constructed the linear model described previously in each subset and then applied that model to SVs in the other set to infer expression effects. We identified significant correlation between the true expression value and the predicted value across all 4 pairwise comparisons (R^2^_CMC_HBCC→CMC_ = 0.4, R^2^_CMC→CMC_HBCC_ = 0.21, R^2^_CMC→CMC_ = 0.41, R^2^_CMC_HBCC→CMC_HBCC_ = 0.22 Figure 4) with deletions (particularly when tested in CMC) consistently performing better.

**Figure 4.**
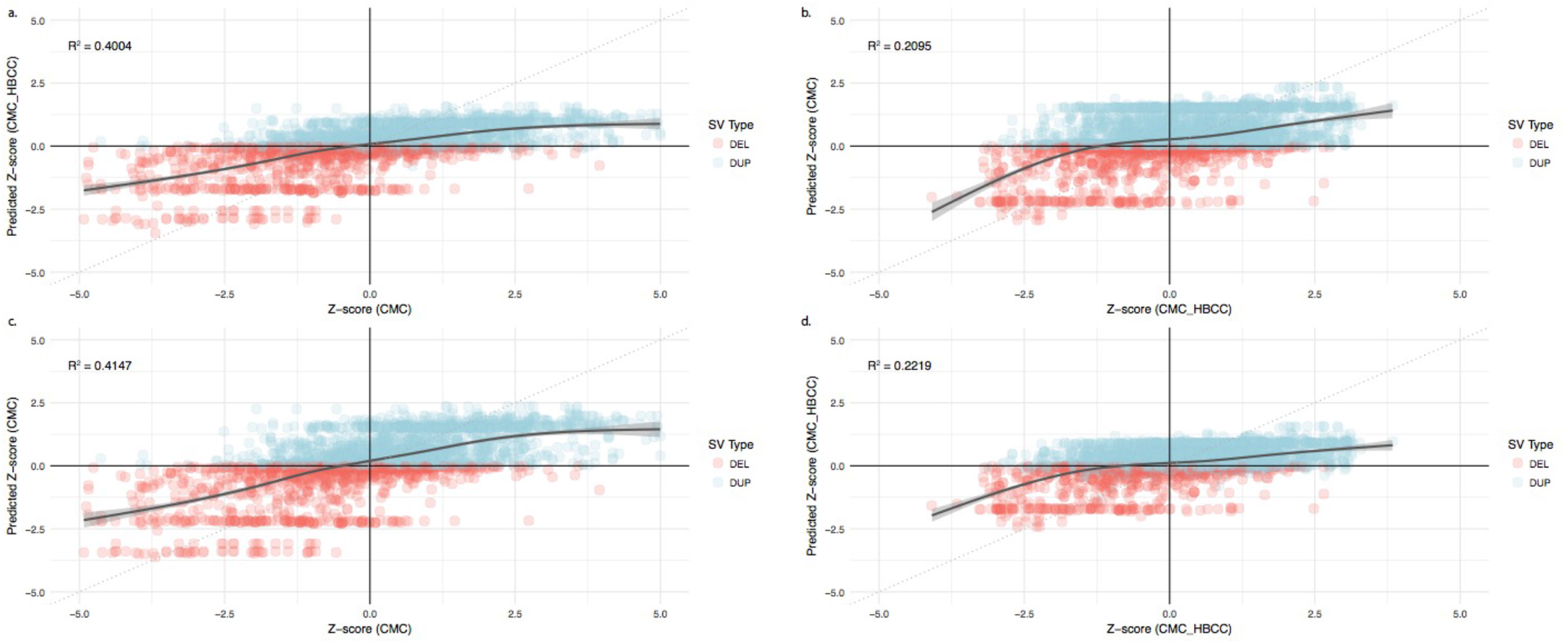
SV expression prediction performance and associated R^2^ building the same linear model using different training and test datasets a) CMC into CMC_HBCC, b) CMC_HBCC into CMC, c) CMC into CMC and d) CMC_HBCC into CMC_HBCC. The best fit line with confidence interval was produced using generalized additive mode smoothing.

Leveraging this model and the previously used measure of genic intolerance to loss-of-function variation, we built an aggregate regulatory disruption score that was the sum of the predicted expression z-scores for each gene weighted by the gene’s intolerance metric (normalized between 0 and 1 with 1 being most intolerant) to annotate SVs. We then applied our model to annotate 98,336 variants in the gnomAD SV dataset^9^ after restricting to CNVs that were below 1% frequency and above 1 kb in length. Of those, 21,476 (21.8%) were predicted to alter the expression of at least one protein-coding gene where we had an intolerance metric and 11,803 of these variants were deletions. We considered deletions in gnomAD as pathogenic if they overlapped at least 50% of any of the 3,455 deletions labeled pathogenic for neurodevelopmental disorders (developmental delay, intellectual disability, or autism) in ClinGen (downloaded from UCSC Genome Browser June 2019). There were 84 variants that met this criterion (39 variants overlapped 100%, as gnomAD includes some individuals with neuropsychiatric disorders). This set of pathogenic deletions had significantly more negative regulatory disruption scores than the rest of the SVs representing a more severe reduction in expression among intolerant genes due to these deletions (p = 3.14×10^−18^, mean score in pathogenic deletions = −5.15, mean score in nonpathogenic deletions = −0.30, Wilcoxon test). Despite ascertainment bias leading to longer deletions being more likely to overlap pathogenic variants, prioritizing variants by regulatory disruption would identify more pathogenic variants than prioritizing by length, with 4 of the top 10 variants being pathogenic if ranked by length (2 complete overlaps) and 6 by regulatory disruption (5 complete overlaps). The regulatory disruption score also better prioritized pathogenic variants than number of genes affected and minor allele frequency, which has limited utility since most deletions (55% or 6,515) have the same frequency, as singletons. (Figure 5, Supplementary Table 3). Overall, regulatory disruption is significantly correlated with whether a deletion is pathogenic (R^2^ = 0.097, p = 1.25×10^−262^). These results indicate the potential of this metric to contribute to improved prioritization of disease causing CNVs.

**Figure 5.**
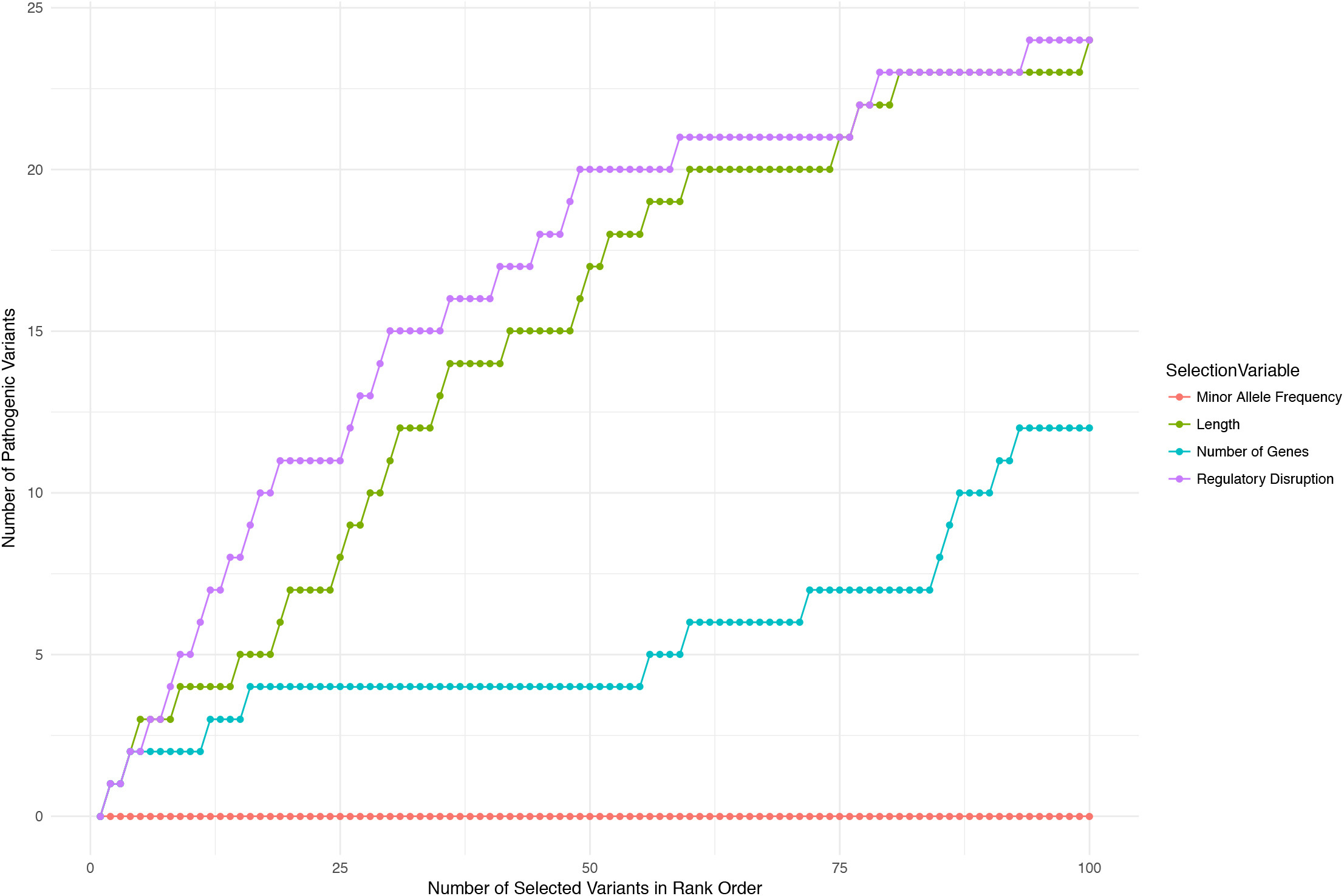
Number of pathogenic variants identified based on rank ordering by length (green), number of genes deleted (blue), minor allele frequency (red) and regulatory disruption (purple). Where multiple variants had the same value, the order was random. Fifty-five percent of deletions (n=6,515) had the lowest frequency (singletons).

## Discussion

The integration of genome and transcriptome data on post-mortem brains from the CMC has provided one of the first opportunities for large-scale characterization of the impact of rare SVs on expression in the brain. Here we demonstrate evidence of selection on rare regulatory SVs, particularly those that alter CTCF binding sites. We found a clear and predictable role for genic and regulatory SVs in altering expression, and we showed that the degree of expression influence is shaped by the intolerance of a gene to deleterious variation. These results suggest the potential to functionally predict and annotate the consequences of SVs on expression. Illustrating this potential, we derived a model to infer expression effects of SVs in independent samples, and applied it to the largest SV resource currently available. This provided evidence that annotating SVs by their regulatory burden could aid in prioritizing disease relevant variants.

Selection maintains deleterious variation at lower allele frequencies, enabling the use of frequency as a proxy for implicating variant classes that may contribute to phenotypes negatively affecting fitness. Here, we showed that SVs overlapping brain regulatory elements including enhancers and CTCF sites were seen at significantly lower frequencies than SVs that were intronic or intergenic. This result remained true even after accounting for SV length and could also be shown in an independent and substantially larger dataset (gnomAD). Regulatory variants, particularly SVs, have the potential to alter the expression of many genes and previous work has already implicated *de novo* variants in fetal enhancers in risk for neurodevelopmental disorders^67^. As an example, a deletion of a CTCF site could enable enhancer activity of nearby genes, a mechanism known as “enhancer hijacking” that is particularly common in cancer^68^. From our data and replication in gnomAD, it appears that SVs altering CTCF sites are as infrequent as genic SVs; this suggests that a substantial proportion may be deleterious and being actively removed from the population by selection. Previous work has directly tested this hypothesis using very different data and approaches and found similar results^64^. With the growing amount of functional and genomic data, assessing the role of SVs on regulatory elements, particularly CTCF sites, in disease will further assess the validity and importance of this finding.

Rare SVs show substantial but variable effects on expression that can be quantified. While identifying potential carriers of rare functional SVs using expression outliers is a practical and valid approach, in our data using this definition would result in the identification of only a small proportion of all genic SVs (including missing most full gene CNVs) and a substantially smaller proportion of regulatory SVs. The approach taken here requires genomic annotations to implicate regulatory effects and an assumption that the annotated elements are functional. Despite these limitations, it is clear that both genic and regulatory SVs have significant effects on dosage that can be quantified and used to infer expression effects of SVs in independent datasets. These effects are strongest when altering coding sequence, but are significant when altering promoter and enhancer sequence as well. In all cases the effect is proportional to the amount of functional sequence affected.

We specifically required that regulatory element annotation be gene-specific to facilitate prediction of enhancer-gene associations. In other words, we showed that we can quantify the effect of a CNV on a specific gene. As ongoing efforts to understand the gene-specific functions of enhancers improve regulatory annotations, so too will the approaches in this paper be improved in accuracy and expanded beyond the roughly 30% of genes we were able to include. Better enhancer-gene target annotations would increase the number of CNVs that could be predicted and the performance of those predictions. We identified a significant negative effect of duplications altering enhancers. This result presents a potentially intriguing implication regarding the direction of regulatory effect of enhancer duplications which may be counter intuitive. One potential concern is that this effect is seen substantially more strongly in one of our two cohorts suggesting a heterogenous or batch effect. Further work is required to better understand the role of enhancer duplications on expression. We did not include CTCF sites in the prediction model as it was not clear how to directly link them to individual genes, however, based on the likely deleteriousness of CTCF SVs, we anticipate effects on potentially many genes and quantifying those effects is an area of future work. We did include TAD annotations as a surrogate for potential contribution to the expression consequence of an SV, and we identify a significant magnification in effect on expression when the SV and gene were within the same TAD, pointing to the importance of TADs in the regulatory landscape. We present a simple linear model that can meaningfully predict expression effects of rare SVs and anticipate that improvements in regulatory annotation and more sophisticated modeling will further the ability to make these predictions.

Improvements in annotation have enabled better prioritization of variants that may cause or increase risk of disease. One avenue to improved annotation has been leveraging large numbers of sequenced individuals in order to quantify the intolerance of each gene to deleterious variation. Here, we show that gene level intolerance metrics also inform regulatory effects of SVs. SVs in intolerant genes have reduced effects on expression, because they are less likely to occur, and when they do occur they alter a smaller proportion of the gene. Combining our SV expression predictions and previously generated gene intolerance measures allowed us to annotate the overall regulatory disruption of an SV by weighting the predicted expression consequences by the relative deleteriousness of the gene affected. For example, a full deletion of a gene where loss-of-function variants are rarely or never seen will be weighted higher than that of a gene that is frequently knocked out in the population. To demonstrate the potential utility of this regulatory disruption score, we annotated all deletions in the gnomAD SV dataset and showed that our annotations were correlated with variants substantially overlapping those that have previously been classified as pathogenic. Further, rank ordering variants by the most extreme regulatory disruption scores enriched for pathogenic variants that would have not been identified by length or frequency. Potential exists for annotation of regulatory disruption to improve prioritization of disease causing or risk increasing SVs; however, further work with clinical or disease samples will be required to fully assess the added value of this approach.

Our work has several limitations, including an assumption that effects of SVs on expression are largely products of proportional alterations on genic and regulatory genomic content. Our predictions demonstrate that this assumption holds on average; however, we anticipate gene and regulatory element specific effects to exist. This assumption was necessary because, in the case of rare SVs, estimating individual variant effects is underpowered. We also leverage assumptions about the direction of effect for large CNVs, and show that this direction of effect extends to smaller variants. We largely focus on CNVs given the expected directional effects as well as the predominance of these variants among our confident calls, which better powers these analyses. We can show expression effects of inversions; however, these effects are substantially smaller and not in a specific direction. We also see very similar patterns of deleteriousness, through lower frequencies, in other classes of SVs that alter regulatory elements and genes. SVs that are not CNVs are still a challenge to call accurately and harder to validate given the smaller set of previously identified variants of high confidence. We anticipate improvements in calling these variants to lead to the ability to better define their functional role in the near future.

This effort was entirely focused on the brain: the expression data were from brain tissues and all of the regulatory elements included were identified in the brain. Some findings, such as the deleteriousness of CTCF sites and the basic prediction model, likely hold true across other tissues as the assumptions are not tissue specific and CTCF sites are relatively shared across tissues. For other results, such as the expression consequences of enhancer SVs or the regulatory disruption scores, it is unclear how well they will generalize given the tissue specific nature of enhancers and the fact that brain expressed genes are among the most conserved and intolerant to variation. It is, therefore, advisable to develop these models in a tissue-specific manner wherever possible.

In conclusion, genome-sequencing and RNA-sequencing when combined in the same samples can be used to interpret the transcriptional consequences of SVs for improved annotation of a variant class that, despite its clear importance, remains difficult to quantify its functional effect and contribution to disease.

## Acknowledgements

This work was supported by R01MH111776 (DMR), R01MH115957, HD081256, and the Simons Foundation for Autism Research Initiative (573206) to MET.. M.L.B. was supported by the National Library of Medicine [T32LM012412]. J.A.C. was supported by the National Institutes of Health [R35GM127087]. The CMC_HBCC genome-sequencing was done through NIH extramural contract HHSN271201400099C. Data were generated as part of the CommonMind Consortium supported by funding from Takeda Pharmaceuticals Company Limited, F. Hoffmann-La Roche Ltd and NIH grants R01MH085542, R01MH093725, P50MH066392, P50MH080405, R01MH097276, RO1-MH-075916, P50M096891, P50MH084053S1, R37MH057881, AG02219, AG05138, MH06692, R01MH110921, R01MH109677, R01MH109897, U01MH103392, and contract HHSN271201300031C through IRP NIMH. Brain tissue for the study was obtained from the following brain bank collections: the Mount Sinai NIH Brain and Tissue Repository, the University of Pennsylvania Alzheimer’s Disease Core Center, the University of Pittsburgh NeuroBioBank and Brain and Tissue Repositories, and the NIMH Human Brain Collection Core. CMC Leadership: Panos Roussos, Joseph Buxbaum, Andrew Chess, Schahram Akbarian, Vahram Haroutunian (Icahn School of Medicine at Mount Sinai), Bernie Devlin, David Lewis (University of Pittsburgh), Raquel Gur, Chang-Gyu Hahn (University of Pennsylvania), Enrico Domenici (University of Trento), Mette A. Peters, Solveig Sieberts (Sage Bionetworks), Thomas Lehner, Stefano Marenco, Barbara K. Lipska (NIMH)

## Author contributions

Conceived and organized the effort: DMR, MT, KJB, MP, PR, BKL, SM. RNA-sequencing generation: TP, GH, PR, JSJ, LS. Structural variant calling: XZ, RLC, HW, HB, MT, DMR. Sample quality control: LH, LS, JSJ. Data analysis and interpretation: DMR, LH, MLB, XZ, JAC, MT, RLC, SKS. Manuscript drafting: DMR, MT, LH, XZ, RLC, SKS. All authors have read and approved of the final version of the manuscript.

