## Supplementary Figures for "Functional annotation of rare structural variation in the human brain"

### Supplementary material

**Supplementary Figure 1.** Counts of SVs by combination of annotations (green) and full set of any annotation (blue)

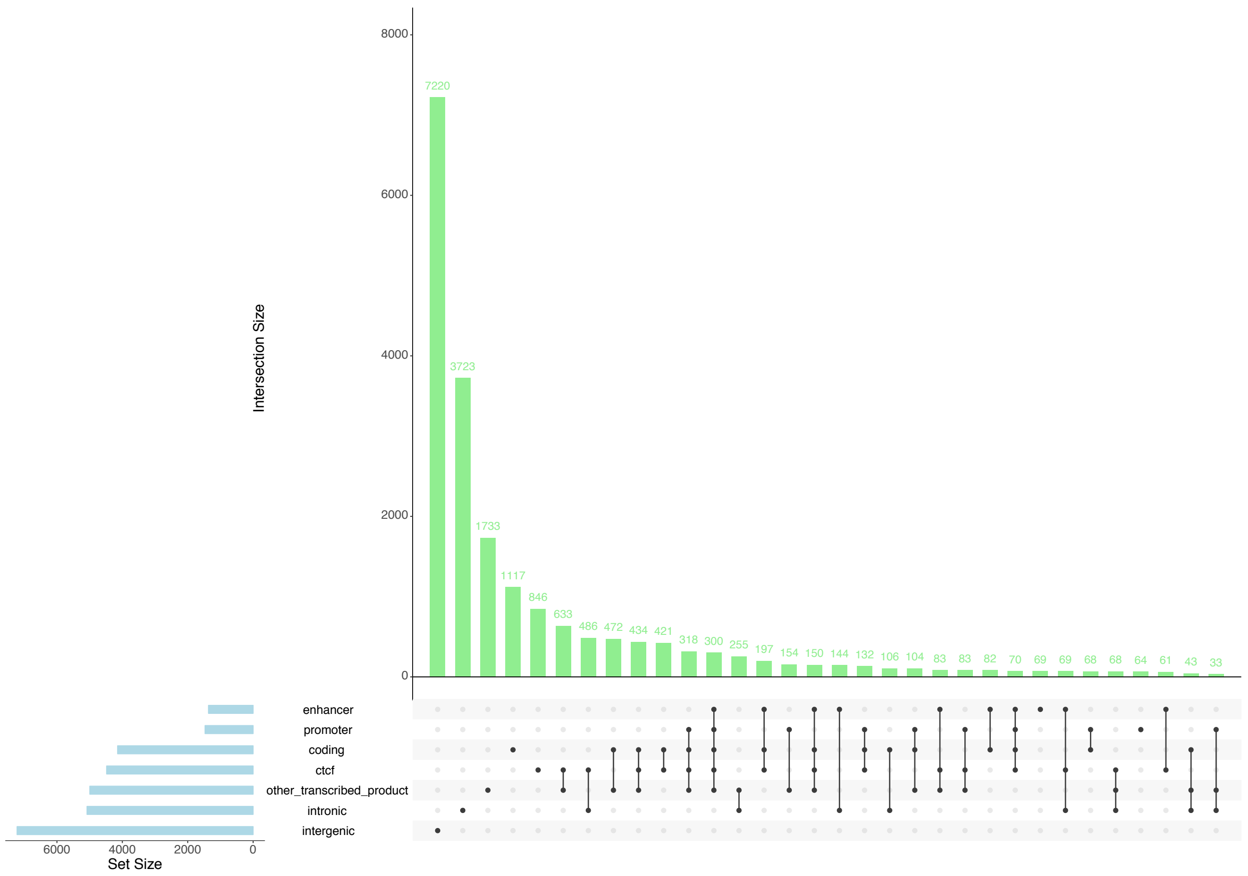

**Supplementary Figure 2.** Mean allele frequency for each of 33 combinations of annotations split by the number of unique annotations and then by SVs that affect all combinations that include CTCF sites, enhancers, coding genes and promoters. Each point represents an annotation combination (or unique annotation). The think bar is the median and the thin bars are 95% confidence interval akin to standard boxplot.

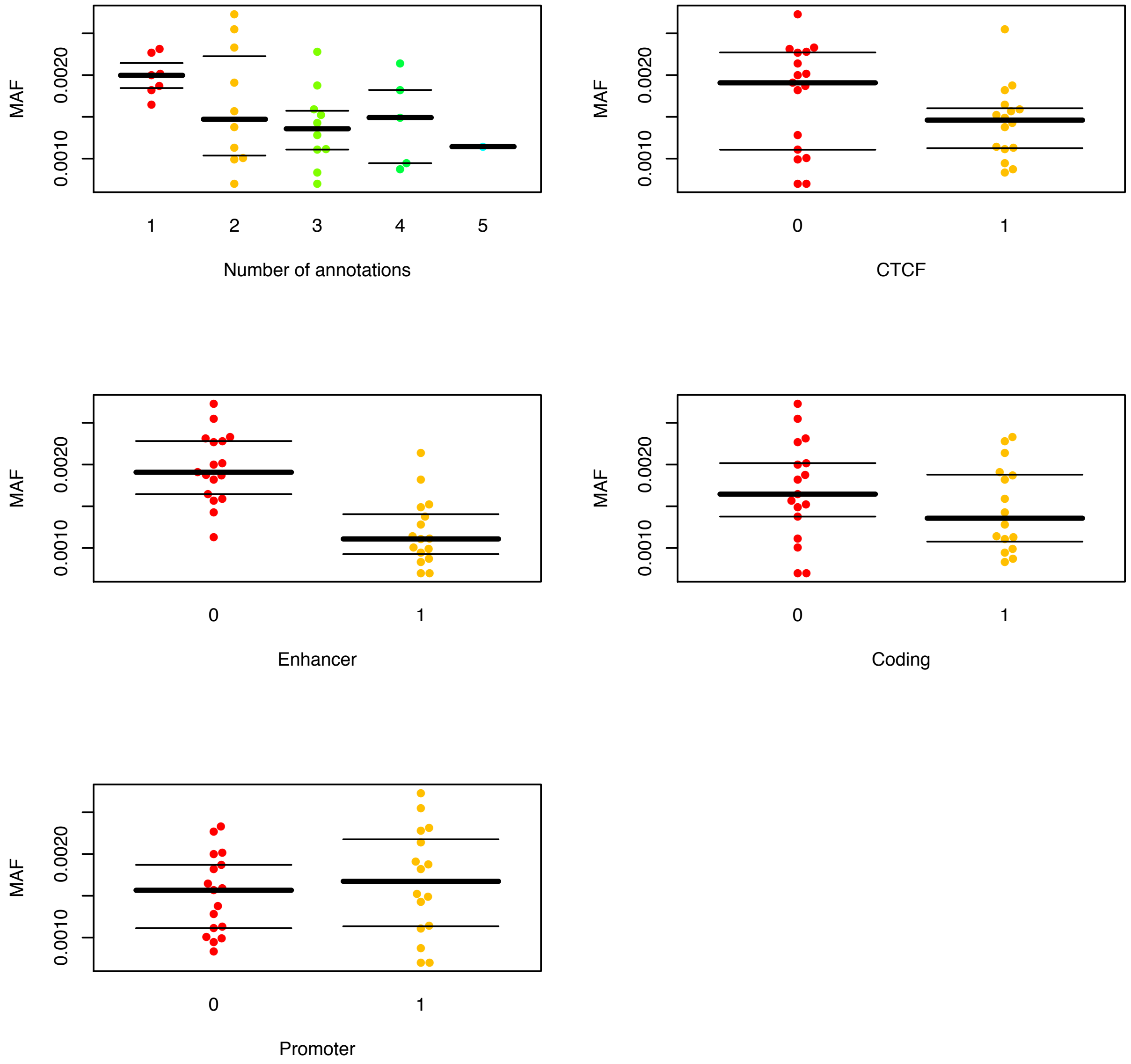

**Supplementary Figure 3.** Distribution plots of the relative expression of full gene deletions or duplications compared to the rest of the CNV in the normalized and covariate adjusted data (normalized) and in the raw counts per million (CPM). Dashed lines represent expected 50% decrease and 50% increase for full gene deletions and duplications respectively.

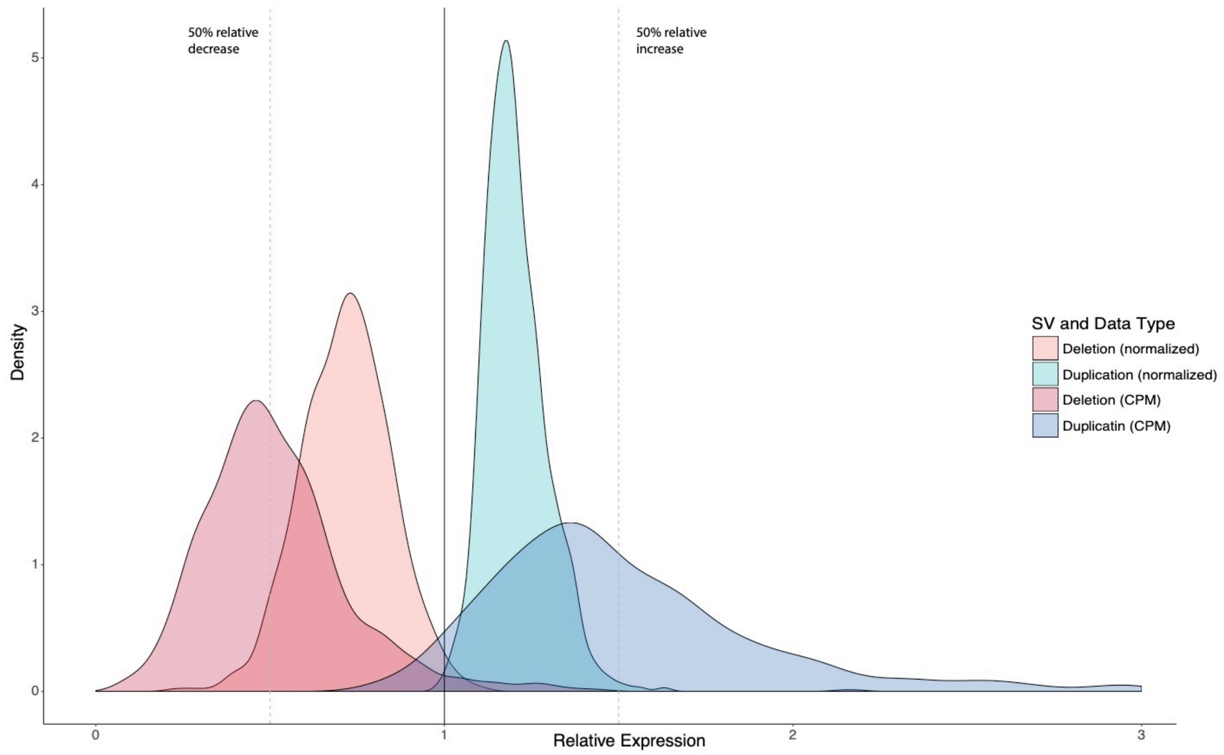

**Supplementary Figure 4.** Correlation of mean expression values per gene/CNV pair between the two independent cohorts (CMC and CMC_HBCC).

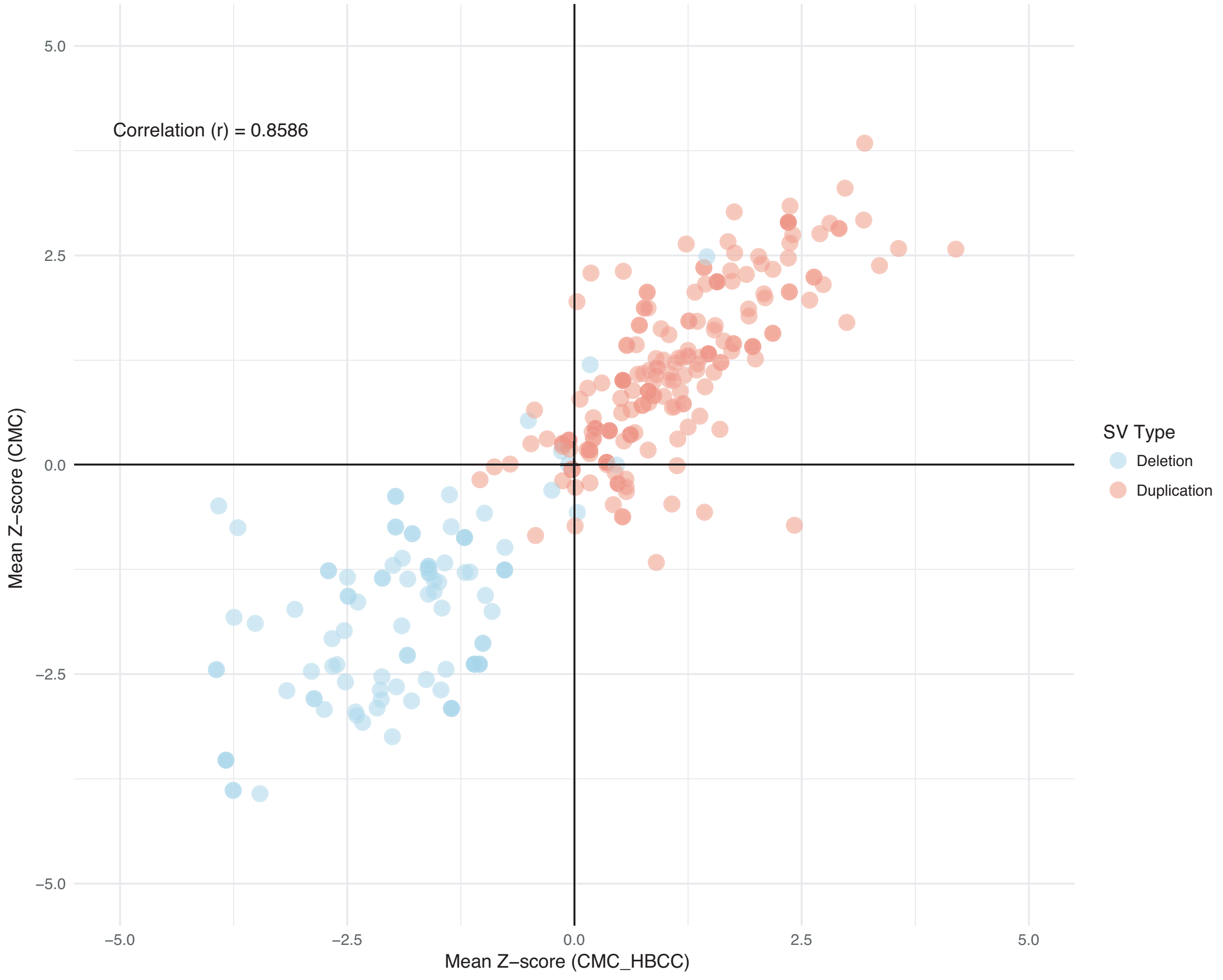

**Supplementary Figure 5.** Distribution of relative expression (Z-score) as a product of the proportion of coding sequence of a gene that is deleted (red) or duplication (blue) stratified by cohort. Lines are loess fits and grey area represents the confidence interval of the line.

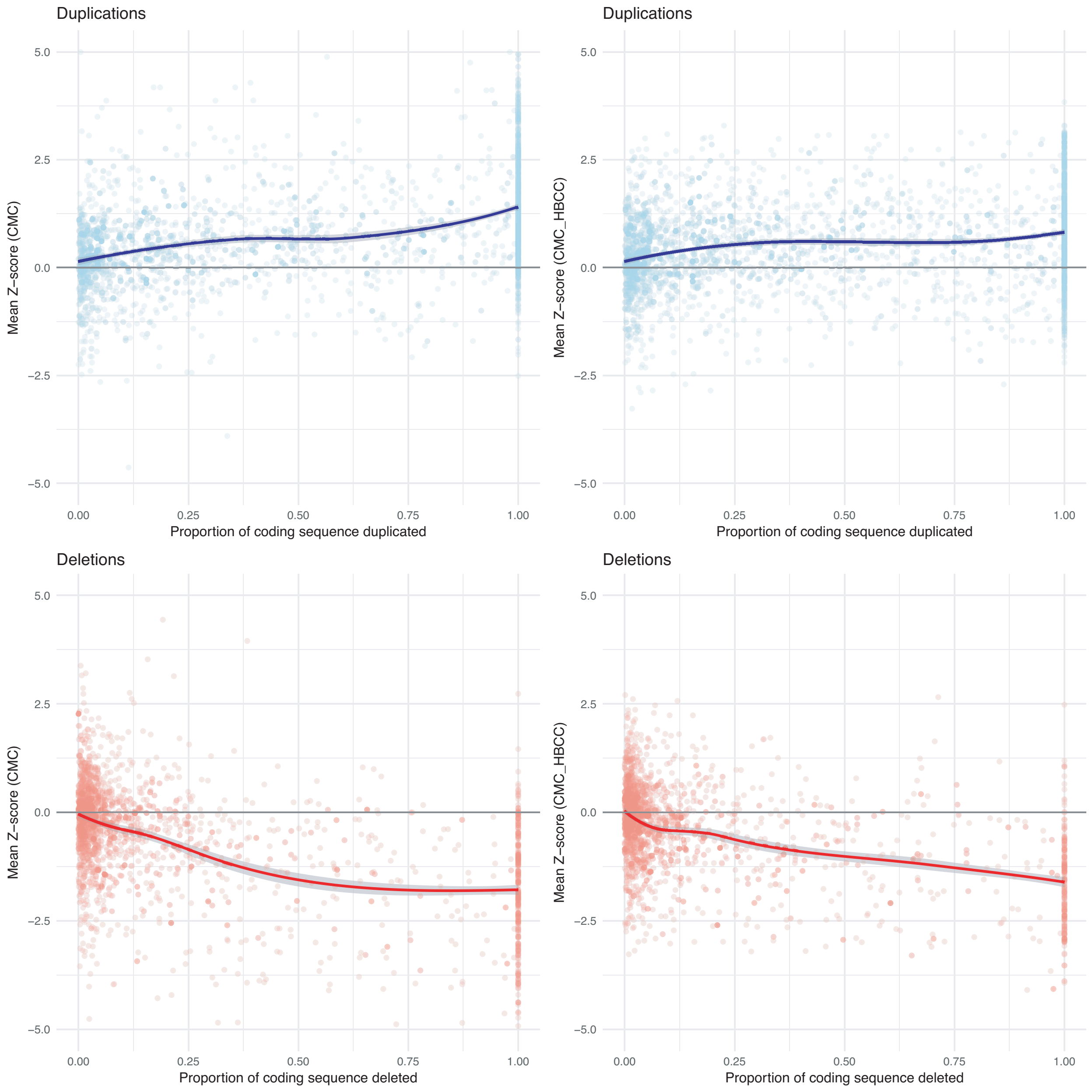
